# A Sign of Disparity: Racial/Ethnic Composition of Treatment Centers is an Independent Risk Factor in SPRINT Trial

**DOI:** 10.1101/102111

**Authors:** Matthew Matlock, Na Le Dang, David Brown, S. Joshua Swamidass

**Affiliations:** Washington University School of Medicine. Saint Louis, MO 63110

**Author notes:** These authors contributed equally (Matthew Matlock, Na Le Dang). This manuscript is a citeable version of a submission to the SPRINT Data Analysis Challenge (https://challenge.nejm.org/pages/home). Except for the addition of an extra figure, this manuscript conforms to the competition word limits of 75 words for the abstract and 700 words for the body of the manuscript.

## Abstract

The racial composition of treatment centers in the SPRINT trial is an independent risk factor for myocardial infarction (MI) and stroke. Independent of individual race or ethnicity, patients face a 39% increase in relative risk of MI or stroke when associated to treatment centers with a high proportion of African Americans. The magnitude of this effect is comparable to smoking. This suggests the strong influence of social determinants on health outcomes among hypertensive patients.

## Background

SPRINT was a randomized, controlled, open-label trial of intensive and standard treatment of hypertension that was conducted at 102 treatment centers in the United States [1]. SPRINT enrolled 30% African Americans, 10% Hispanics and 58% whites. The racial and ethnic composition of the treatment centers (RC) may be a surrogate marker for social determinants of disease, such as poverty, health care access, nutritional access, and health care literacy. We hypothesize that center-level RC will influence the outcomes of patients enrolled in the SPRINT trial, even after controlling for other factors, including the race of individual patients.

## Methods

We used logistic regression to study the relationship between RC and myocardial infarction (MI) and stroke (the primary outcome) in the SPRINT population. To the clinical and demographic data, we added the RC of the center at which each patient was treated. The model was trained to predict adverse outcome during the study period. The model achieved moderate accuracy (65.7% Area Under the ROC Curve), which is not accurate enough to triage individual patients. However, it is accurate enough to determine the importance of individual factors.

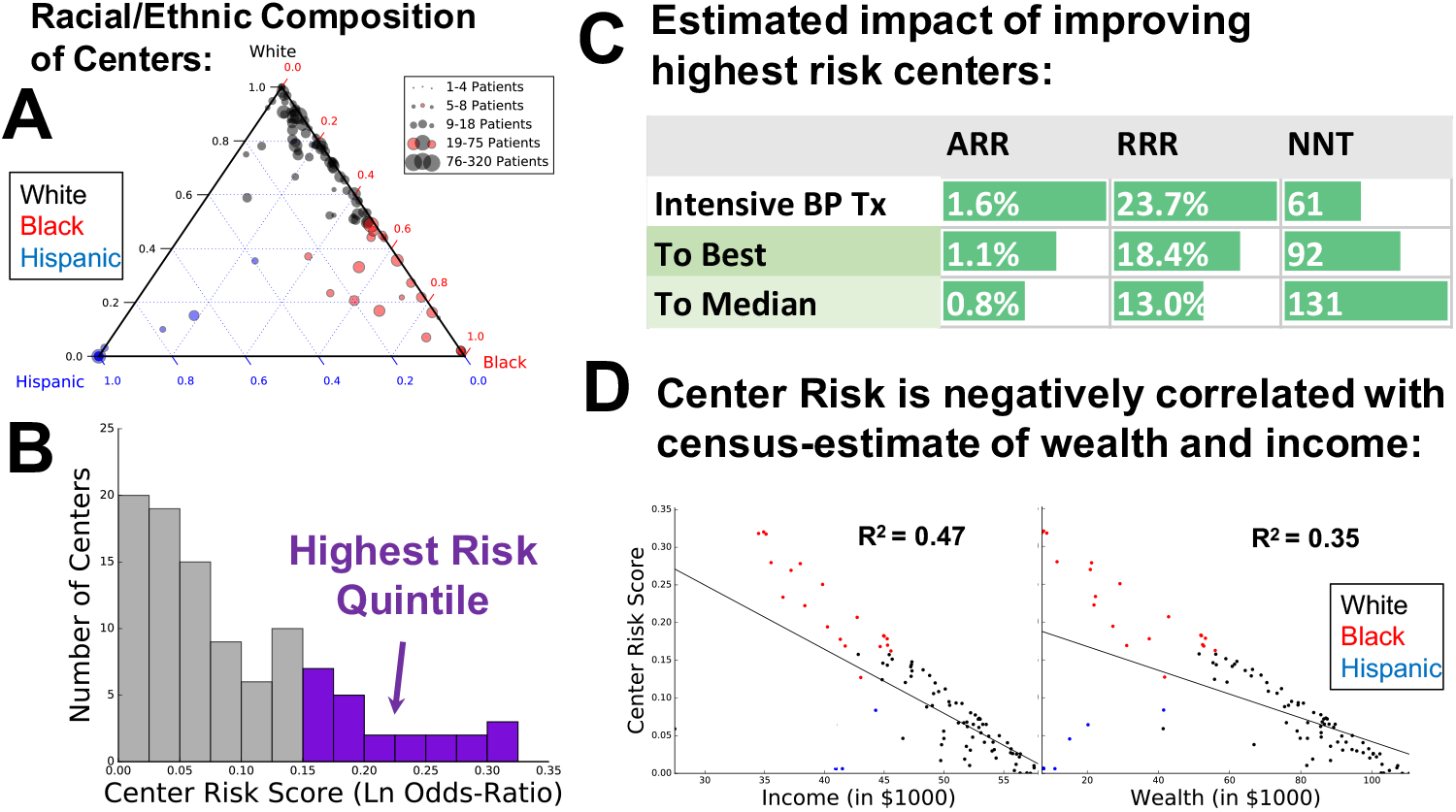

## Results

First, the 102 treatment centers in the SPRINT trial represent a wide range of racial/ethnic composition, from predominantly White to Black or Hispanic (Panel A). Nonetheless, 1,848 patients (430 White/Other, 1,149 Black and 269 Hispanic) were treated in a center where they were a minority. Critically, because of these cross over patients, the effect of center-level RC can be tested, while controlling for the effect of race at an individual level.

Second, we studied the regression coefficients to understand the importance of center-level RC. In this test, individual race/ethnicity are not statistically significant: Black OR 0.90 (P=0.17) and Hispanic OR (P=0.19). At the center level, Hispanic composition is not significant (OR 1.01, P=0.97). However, Black composition is a significant factor with a strong magnitude of effect (OR 1.38, 95% CI: 1.11 - 1.73, P=0.0045).

Third, the magnitude of effect of center RC is clinically significant. This effect is comparable to (but opposite) the effect of intensive treatment for BP (OR 0.74), and essentially equal to having a history of smoking (OR 1.33). Studying this further, the model’s coefficients were used to compute a center level risk score, which ranged from 0.00 to 0.32 log-odds (Panel B); so, the highest risk centers increase relative risk by 39%.

Fourth, we simulated the MI and stroke outcomes achieved by modifying center-level risk. This simulates interventions that successfully identify and rectify the root causes for the increased risk among patients in the high risk centers. This might correspond with moving patients to lower risk centers, or to alleviating specific social determinants of disease. We reduced the center risk of the 1872 patients in the highest risk quintile of centers (purple bars in Panel B) to that of the best and the median center. Improving their center risk improved outcomes substantially (Panel C): to best risk (RRR 18.4%, NNT 92) and median risk (RRR 13.0%, NNT 131). This effect was even stronger when limiting the simulation to the 381 patients in the five highest risk centers: to best risk (RRR 24.9%, NNT 70) and median risk (RRR 19.9%, NNT 88). The effect on outcome is comparable to the benefit of intensive treatment for hypertension (RRR 23.7%, NNT 61).

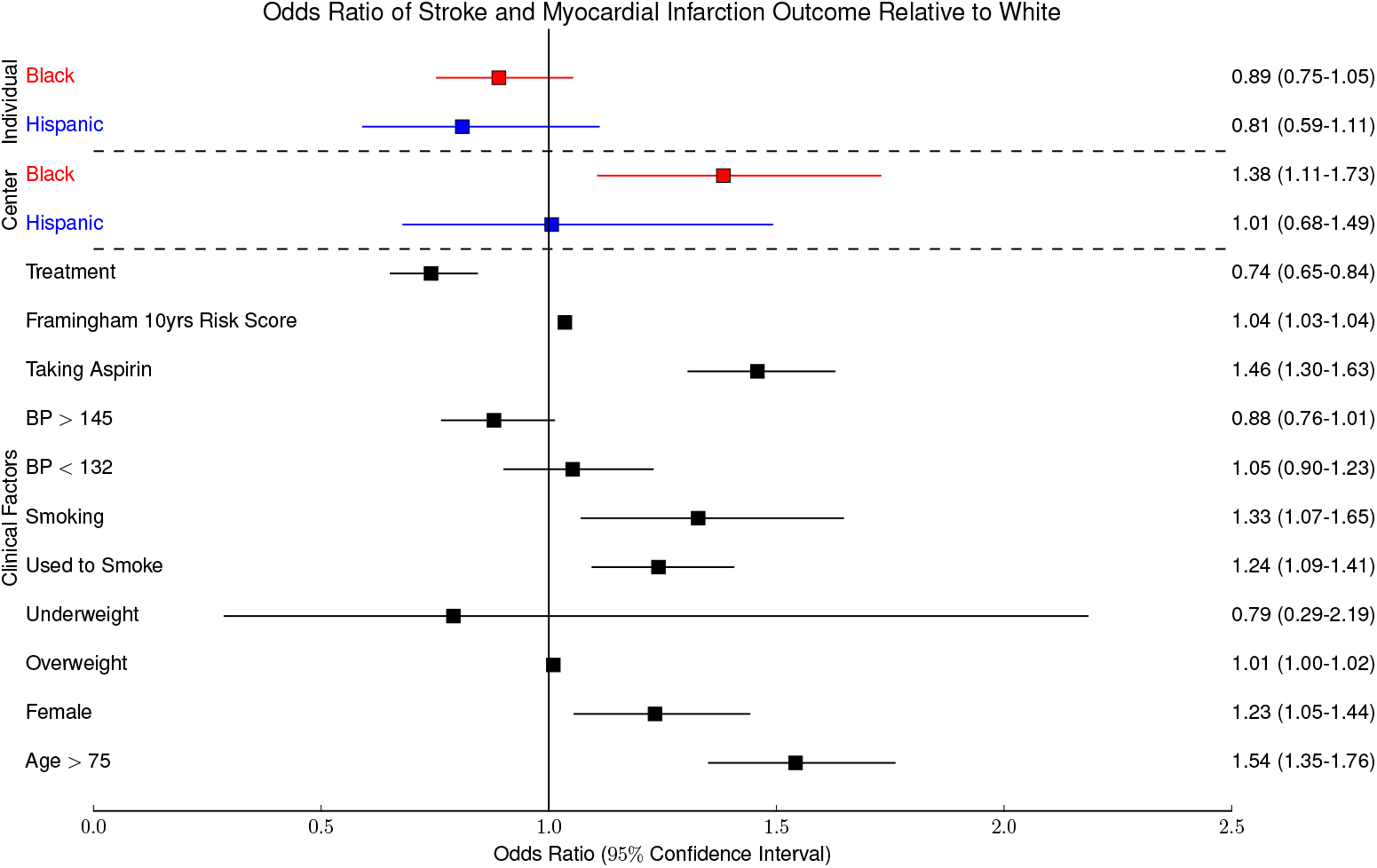

## Conclusion

This analysis encourages inquiry into the root mechanisms of this effect.

It is possible that center-level RC is a surrogate for systemic factors external to the clinic. For example, RC could indicate the poverty of a patient population, and that poverty may be driving the association with poor outcomes. We expect center-level risk is closely correlated with the socio-economic status of patients; a census-based estimate of income and wealth confirms this (Panel D). [2] However, more factors are required to explain the discrepancy between Hispanic and Black RC (no effect vs. strong effect).

Identifying the mechanisms that give rise to center-level risk might suggest effective interventions. These interventions are important, and might improve health for all of us.

## Acknowledgments

We thank the New England Journal of Medicine and the SPRINT research team for conducting the Data Analysis Challenge. Research reported in this publication was supported by the National Library Of Medicine of the National Institutes of Health under award numbers R01LM012222 and R01LM012482. We also thank both the Department of Immunology and Pathology at the Washington University School of Medicine and the Washington University Center for Biological Systems Engineering for their generous support of this work.

